# Prediction of evolutionary constraint by genomic annotations improves prioritization of causal variants in maize

**DOI:** 10.1101/2021.09.03.458856

**Authors:** Guillaume P. Ramstein, Edward S. Buckler

## Abstract

Crop improvement through cross-population genomic prediction and genome editing requires identification of causal variants at single-site resolution. Most genetic mapping studies have generally lacked such resolution. In contrast, evolutionary approaches can detect genetic effects at high resolution, but they are limited by shifting selection, missing data, and low depth of multiple-sequence alignments. Here we used genomic annotations to accurately predict nucleotide conservation across Angiosperms, as a proxy for fitness effect of mutations. Using only sequence analysis, we annotated non-synonymous mutations in 25,824 maize gene models, with information from bioinformatics (SIFT scores, GC content, transposon insertion, k-mer frequency) and deep learning (predicted effects of polymorphisms on protein representations by UniRep). Our predictions were validated by experimental information: within-species conservation, chromatin accessibility, gene expression and gene ontology enrichment. Importantly, they also improved genomic prediction for fitness-related traits (grain yield) in elite maize panels (+5% and +38% prediction accuracy within and across panels, respectively), by stringent prioritization of ≤ 1% of single-site variants (e.g., 104 sites and approximately 15deleterious alleles per haploid genome). Our results suggest that predicting nucleotide conservation across Angiosperms may effectively prioritize sites most likely to impact fitness-related traits in crops, without being limited by shifting selection, missing data, and low depth of multiple-sequence alignments. Our approach – Prediction of mutation Impact by Calibrated Nucleotide Conservation (PICNC) – could be useful to select polymorphisms for accurate genomic prediction, and candidate mutations for efficient base editing.

## Background

In quantitative genetics, causal mutations are generally detected by statistical associations between genetic polymorphisms and phenotypic differences within species (QTL effects). QTL effects are useful in plant breeding (e.g., in genomic prediction), but they may be confounded by the co-segregation of neutral polymorphisms with causal mutations (linkage disequilibrium; LD) [1]. In contrast, phylogenetic nucleotide conservation (PNC) detects causal mutations by conservation of DNA bases across species. This statistic is an indirect indicator of fitness effect [2], but it is less confounded by LD, due to the uncoupling of causal mutations and nearby polymorphisms at large evolutionary timescales. PNC, as quantified by methods like SIFT [3] or gerp++ [4], may support plant breeding techniques which require identification of causal mutations at single-site resolution (e.g., cross-population genomic prediction, CRISPR-based editing).

Despite key advantages, PNC has practical disadvantages which limit its usefulness in quantitative genetics [5, 6]: (*i*) it is calculated from a multiple-sequence alignment (MSA), which requires cross-species conservation of alignable genomic regions; (*ii*) it can be so sensitive that maximum constraint can be reached even at moderate fitness effects, due to the exponential relationship between fitness effects and fixation probability of mutations [2, 7]; and (*iii*) it may be biased by functional turnover (shifting selection) and clade-specific conservation. To overcome these limitations, PNC may be predicted throughout the genome, based on annotations which capture the genomic characteristics of fitness effects (genomic annotations). Previous methods like CADD [8, 9] and LINSIGHT [10, 11] have been introduced to predict PNC. However, they have relied on genomic annotations from large-scale experiments in human, which may not be available in plants. Moreover, the spatial resolution of their inference has been limited by small evolutionary timescales, within human and across related species.

In this study, we introduce a machine learning method to predict PNC across Angiosperms in coding regions in maize (*Zea mays* L.), using genomic annotations that are readily available from DNA and protein sequence data. Computational annotations have several advantages: low cost, absence of missing values, and ease of portability from one genome to another. They may also provide latent (non-observed) representations of genes, and can be used to perform *in silico* mutagenesis to predict the impact of point mutations on these representations. To achieve high resolution and high accuracy, we predict PNC at large evolutionary timescales by high-resolution genomic annotations. We use *in silico* mutagenesis to estimate the effect of mutations on protein structure, based on UniRep, a sequence-based deep learning technique which characterizes protein structure by latent representations of protein sequences [12]. Our predictions of PNC are validated by functional enrichment. Importantly, our validations include cross-population genomic prediction, in which genome-wide single-nucleotide polymorphisms (SNPs) are used to predict agronomic traits, and SNPs in coding regions are upweighted according to their predicted PNC. Together, our functional analyses show that predicted PNC is useful to identify impactful genes and SNPs for fitness-related traits in maize.

## Results

### Monomorphic sites in maize are under stronger evolutionary constraint than polymorphic sites

In this study, we aimed at capturing the genomic basis for fitness effects in coding regions in maize, by predicting PNC at nonsynonymous mutations from genomic annotations. The DNA bases (genomic sites) with large fitness effects are subjected to evolutionary constraint (negative selection), so they should be conserved across species, and monomorphic within species. Accordingly, monomorphic sites within maize tended to be more conserved across Angiosperms, compared to SNPs: they were aligned in MSAs over larger evolutionary timescales (Supplementary Fig. S1), and their ‘rejected substitutions’ (RS) scores were higher (Fig. 1).

**Figure 1.**
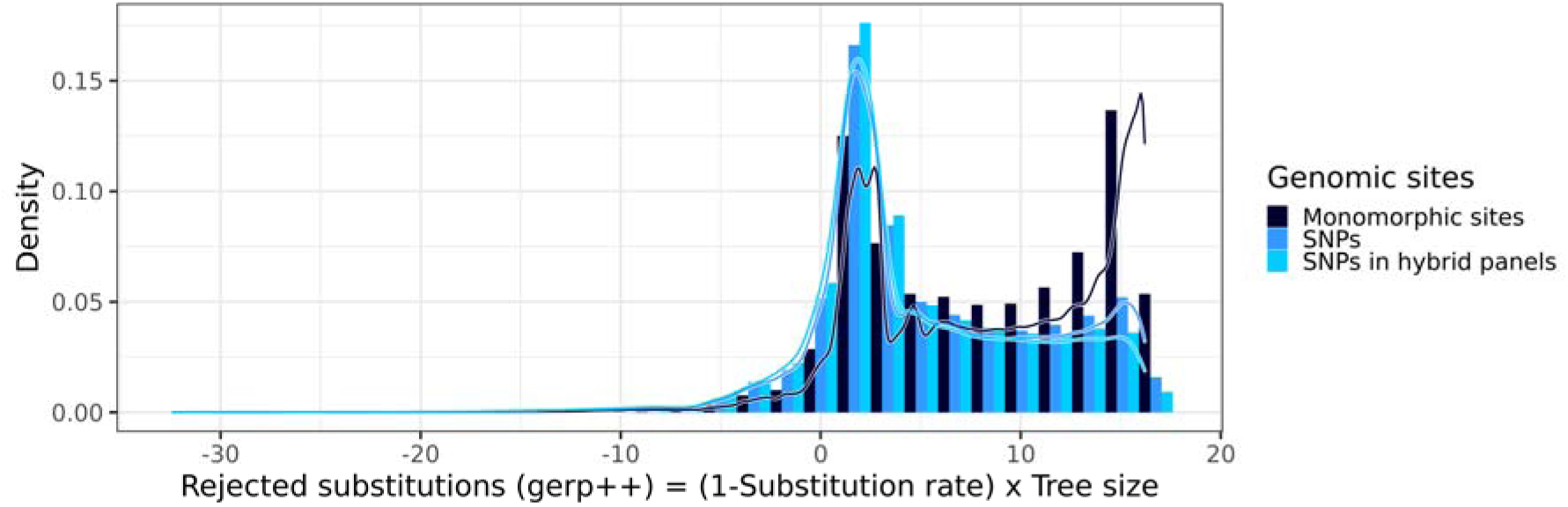
Distribution of rejected substitution (RS) scores by category of DNA bases. RS scores, which integrate information about conservation (1-Substitution rate) and MSA depth (Tree size), were calculated by gerp++ [4] as previously described [23]. Monomorphic sites: sites with no observed polymorphism within maize. SNPs: observed polymorphisms in Hapmap 3.2.1, a representative panel of inbred lines in maize [13]. SNPs in hybrid panels: subset of SNPs which are also observed in two panels of hybrid crosses between inbred lines and testers [14].

Our approach – Prediction of mutation Impact by Calibrated Nucleotide Conservation (PICNC) – used conserved sites as positive examples for large fitness effects, and sites in non-aligned regions as negative examples for neutral effects. PICNC did not rely on within-species variability, so we could include nonsynonymous mutations at monomorphic sites in maize for model training. This helped us avoid survivorship bias at SNP sites and provide many more instances of PNC to learn about the genomic characteristics of fitness effects: 20,136,310 monomorphic sites instead of 483,448 SNPs in Hapmap 3.2.1 (the reference panel in maize) [13] or 103,905 SNPs in elite maize panels (hybrid panels) [14] (Fig. 1, Supplementary Fig. S1).

### Evolutionary constraint is accurately predicted by genomic annotations from sequence analysis

At each nonsynonymous mutation, PNC was characterized by a deep MSA (tree size > 5 expected nucleotide substitutions under a neutral model) and a high nucleotide conservation (substitution rate < 0.05 in the MSA at the site of the mutation). Observed PNC was used to train a probability random forest with genomic annotations about genomic structure (transposon insertion, GC content, average *k*-mer frequency) and protein structure (SIFT score, mutation type, protein features and *in silico* mutagenesis scores from UniRep). Our prediction approach benefited from three key advantages (Fig. 2): (*i*) monomorphic sites provided more information about PNC; (*ii*) annotations like SIFT scores and *in silico* mutagenesis scores from UniRep enabled predictions at single-site resolution; and (*iii*) leave-one-chromosome-out prediction avoided overfitting to observed PNC.

**Figure 2.**
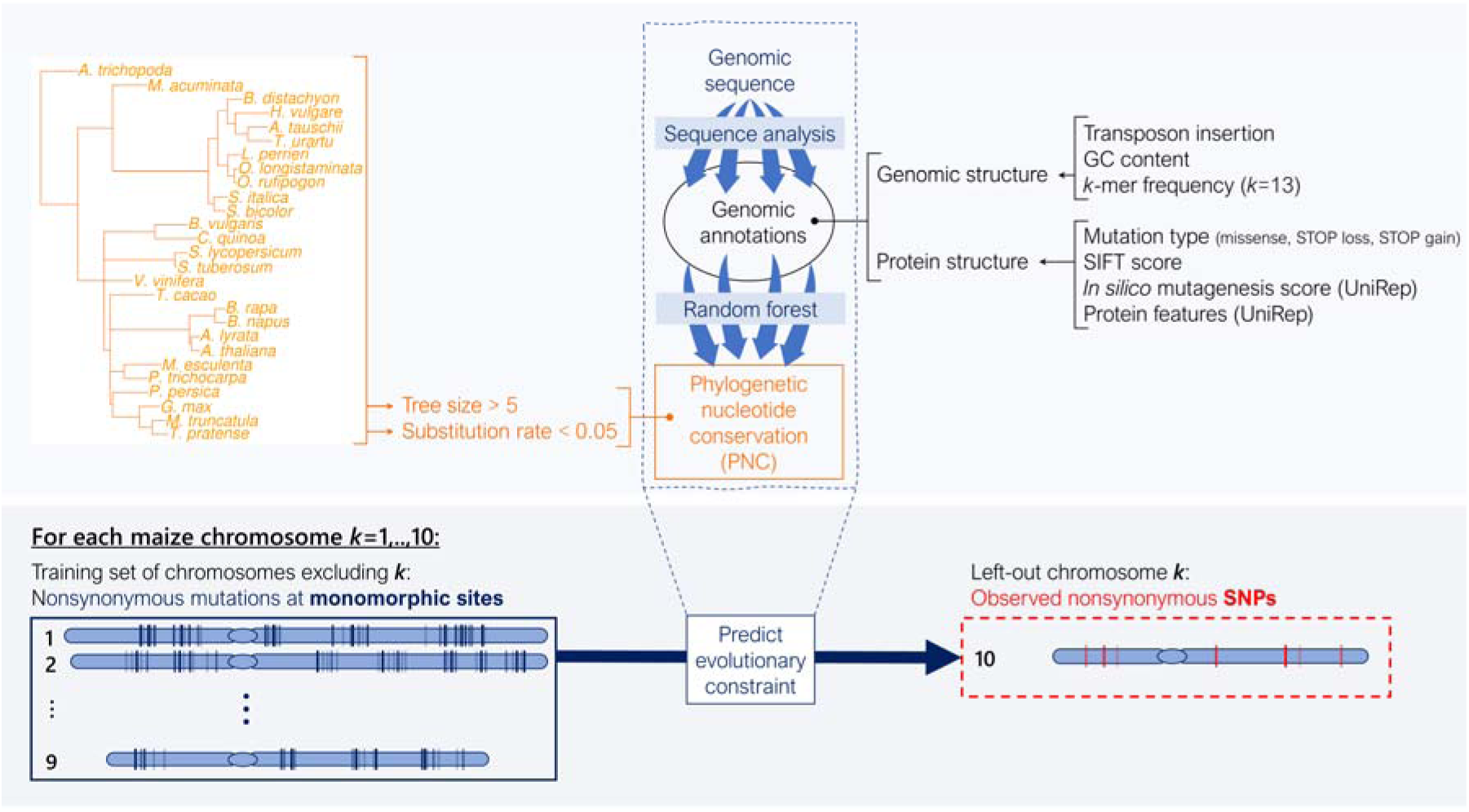
Prediction of mutation Impact by Calibrated Nucleotide Conservation (PICNC). Methodology for prediction of phylogenetic nucleotide conservation (PNC) by probability random forests. PNC was defined by high conservation (substitution rate < 0.05) over deep MSA (tree size > 5 expected neutral substitutions). Genomic annotations were produced only by sequence analysis. They described genomic structure and protein structure at nonsynonymous point mutations in maize coding regions. Monomorphic sites (no observed polymorphism within maize) were used for training, and observed SNPs were used for prediction. In leave-one-chromosome-out prediction, a probability random forest is trained ten times, once for each left-out chromosome.

Compared to a baseline model including SIFT score and mutation type (missense, stop gain, stop loss), annotations about genomic structure (especially GC content) contributed to an improved prediction accuracy for PNC, from 72% to 76% (Fig. 3a,b). Protein features (UniRep variables) and their *in silico* mutagenesis scores resulted in a further increase to > 80% (Fig. 3a). This additional gain in accuracy suggests that novel annotations about protein structure and the impact of nonsynonymous mutations may improve our ability to detect deleterious mutations. As expected, SIFT score was the most useful genomic annotation for predicting PNC, but its importance was on par with those of UniRep variables and their *in silico* mutagenesis scores (Fig. 3b).

**Figure 3.**
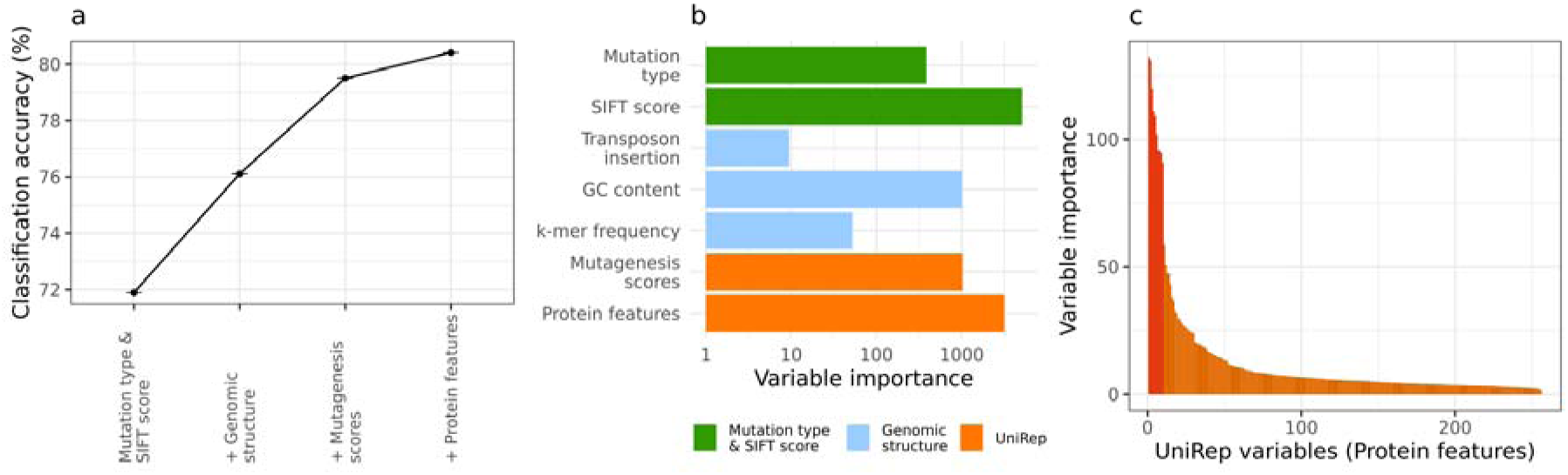
Contribution of genomic annotations to prediction accuracy in probability random forests. (a) Classification accuracy of probability random forests for predicted phylogenetic nucleotide conservation (PNC). Accuracy: percentage of correct calls, i.e., the percentage of sites for which predicted PNC (rounded) equaled observed PNC, over three replicates. Accuracy was weighted to account for imbalance with respect to PNC and chromosome (see Methods). Sets of genomic annotations were sequentially added to the set of predictors in probability random forests. Mutation type & SIFT score: Mutation type (missense, stop gain or stop loss), SIFT score (with missing values set to 1) and SIFT class (“constrained” if SIFT score ≤ 0.05, “tolerated” otherwise). Genomic structure: GC content, *k*-mer frequency and transposon insertion. Mutagenesis scores: *in silico* mutagenesis scores for UniRep variables. Protein features: UniRep variables, generated by the 256-unit UniRep model. (b) Variable importance of genomic annotations. Variable importance: corrected impurity measure in probability random forests [59]. (c) A subset of 10 UniRep variables stood out as contributing most to the prediction accuracy for PNC.

To investigate the usefulness of UniRep variables within maize, we fitted random forests models which regressed gene properties on UniRep variables. UniRep variables captured gene variability within maize, for expression levels (RNA and protein abundance) and selective constraint (negatively associated with the nonsynonymous-to-synonymous SNP ratio, P_n_/P_s_) (Pearson correlation > 0.35; Supplementary Fig. S2). Therefore, the UniRep variables, which were designed to capture protein structural variability across viruses, prokaryotes and eukaryotes, were useful, both across Angiosperms (on PNC) and within maize (on gene properties). Interestingly, a subset of 10 variables stood out as capturing more information about PNC (Fig. 3c), and was also important for predicting selective constraint within species, as reflected by P_n_/P_s_ (Supplementary Fig. S3) [15]. Therefore, few UniRep variables may capture the fitness effects of maize genes, and could serve as succinct functional representations of genes for effects on fitness-related traits.

### Predicted evolutionary constraint identifies deleterious variants in maize

Observed PNC is prone to errors and lacks power to discriminate among different sizes of fitness effects [15]. On the other hand, predicted PNC is estimated by smooth functions of genomic annotations learned across many sites. Therefore, we tested the hypothesis that predicted PNC could estimate fitness effects more accurately than observed PNC. Variability at SNPs, as reflected by minor allele frequency in Hapmap 3.2.1 (MAF), provided information about selective constraint at DNA sites within species. The relationship between PNC and fitness effects was corroborated by its negative association with MAF, as was previously reported [16]. Notably, SNPs prioritized by predicted PNC tended to have lower MAF as prioritizations grew more stringent, and these SNPs were eventually much rarer than those prioritized by observed PNC (Fig. 4a, Supplementary Fig. S4). The functional relevance of predicted PNC was also supported by its positive association with chromatin accessibility (Fig. 4b), which is correlated with phenotypic effects in maize [14, 17]. However, there was a significant increase in expression QTL (eQTL) effect only for observed PNC (P = 0.003 and P = 0.034 in shoot and root tissues respectively, compared to P = 0.120 and P = 0.485 for predicted PNC; Fig. 4c), possibly because the genomic annotations used to predict PNC lack relevant information about gene expression, or because the sites in eQTL are not under strong negative selection across Angiosperms.

**Figure 4.**
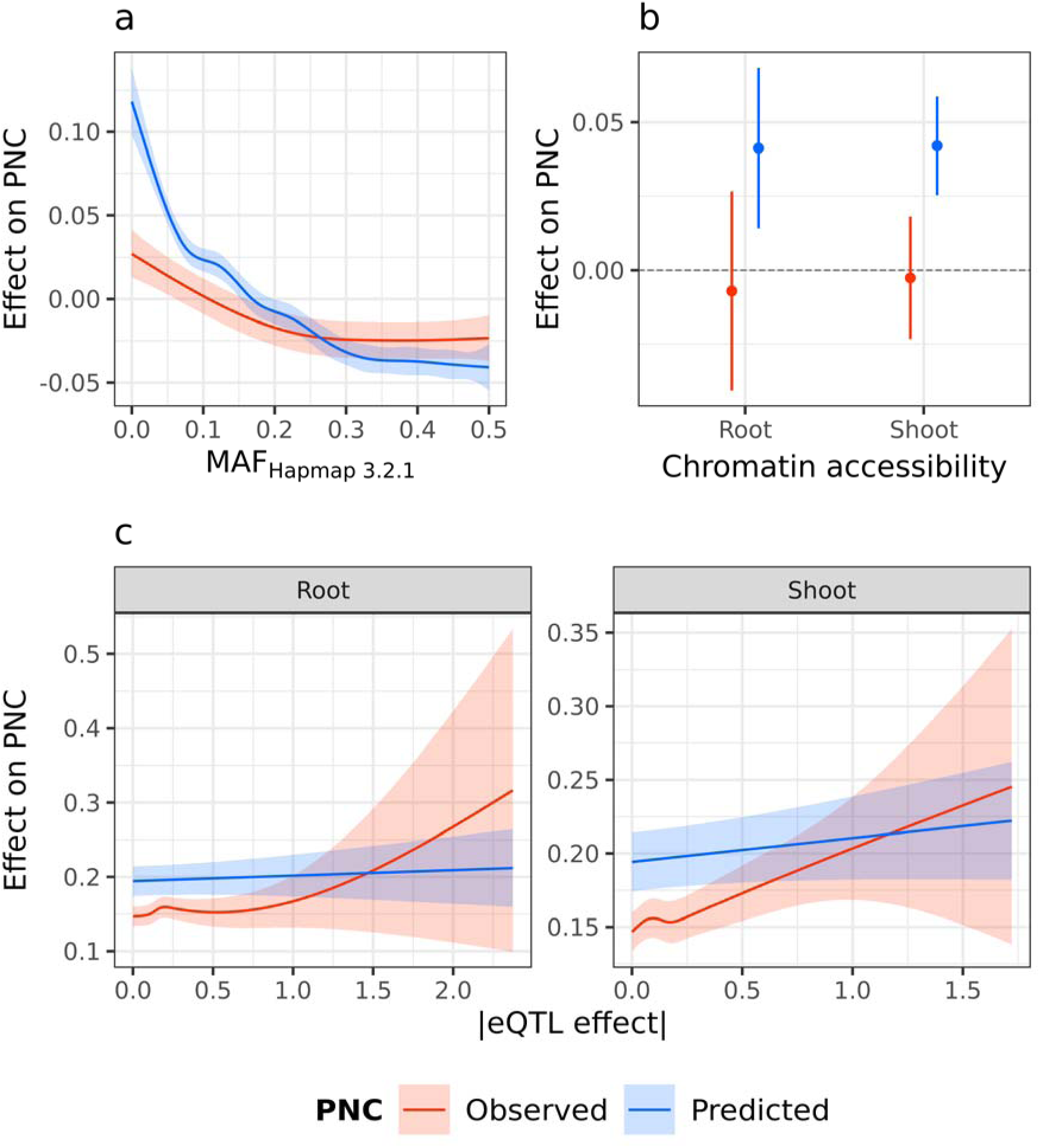
Relationship between phylogenetic nucleotide conservation (PNC) and experimental annotations at SNPs. (a) Decrease in observed and predicted PNC over within-species variability, quantified by MAF in Hapmap 3.2.1 [13]. (b) Increase in predicted PNC in accessible chromatin regions, defined by MNase hypersensitivity in shoot or root tissues [17]. (c) Positive association between observed PNC and expression QTL effect (in absolute values) in shoot or root tissues, estimated in a diverse maize panel [61].

### Predicted evolutionary prioritizes functional genes in maize

Under the hypothesis that predicted PNC identifies impactful genes, the set of genes prioritized by predicted PNC should be enriched for important functional attributes like high gene expression. Observed PNC resulted in significant enrichment for highly-expressed genes (higher RNA and protein abundance, in more tissues), among 14,646 prioritized genes out of the 24,549 genes containing nonsynonymous SNPs. However, such enrichment was more evident with predicted PNC, and increased consistently as fewer genes were selected (Fig. 5a). As expected, P_n_/P_s_ also decreased consistently (Supplementary Fig. S5). These results suggest that predicted PNC pointed to impactful genes. Alternatively, PNC at these prioritized genes may be a direct consequence of “expression-rate anticorrelation”, i.e., selection against cytotoxic byproducts of highly expressed genes (e.g., due to mRNA misfolding or protein misinteraction), rather than direct selection for their functional importance [18–22].

**Figure 5.**
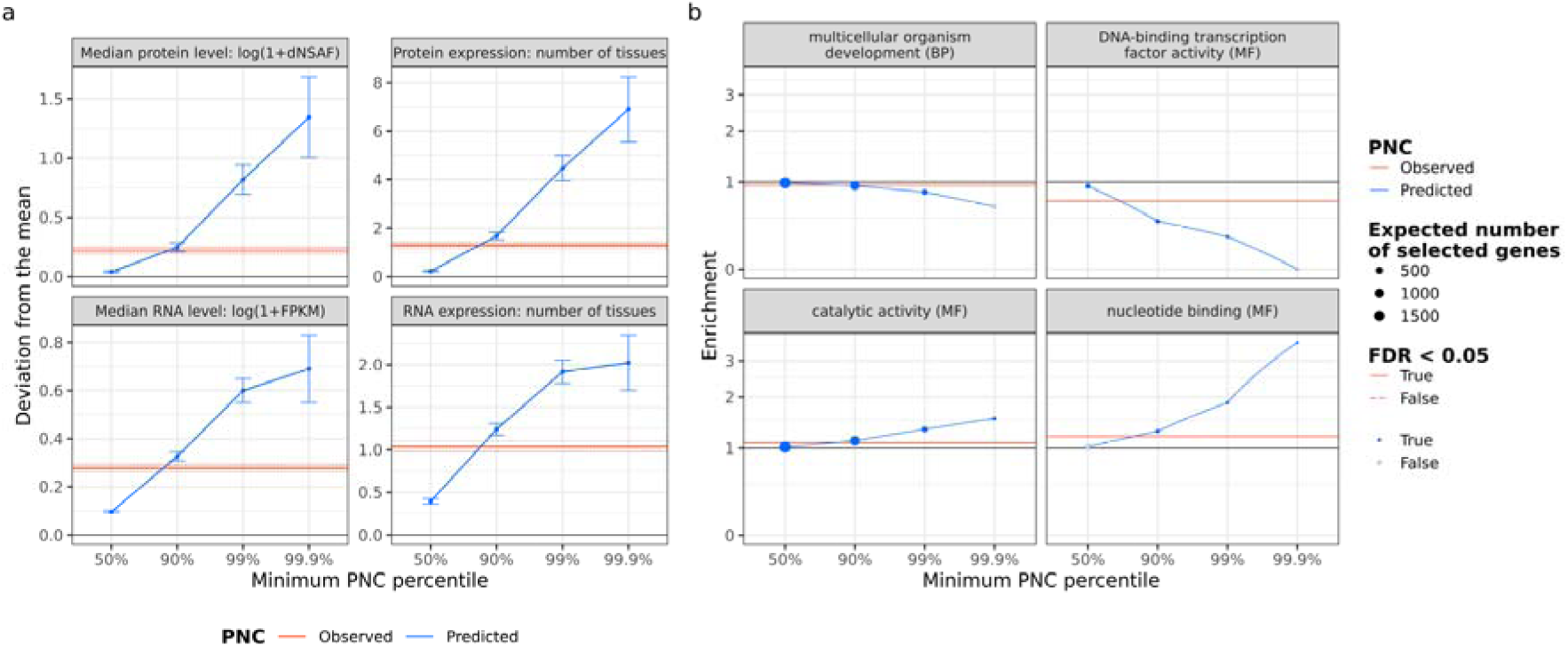
Functional enrichment of genes prioritized by phylogenetic nucleotide conservation (PNC). Genes were prioritized by selecting SNPs with a predicted PNC above the 50%, 90%, 99%, or 99.9% quantile, or observed PNC equal to 1 (tree size > 5, substitution rate < 0.05). (a) Difference in average expression between prioritized genes and all genes. Gene expression is quantified by RNA abundance (FPKM over 23 tissues) and protein abundance (dNSAF over 32 tissues) based on the gene expression atlas of [64]: median expression, and number of tissues with non-zero expression level. Error bars and dotted lines represent 95% confidence intervals in two-sample t-tests, for predicted and observed PNC respectively. (b) Enrichment of prioritized genes for gene ontology (GO) classes. Ratio of number of prioritized genes over expected number under the null hypothesis (random gene prioritization). GO classes belong to the plant GO slim subset. Ontology: BP, biological process; MF: molecular function. For each threshold and ontology, false discovery rates (FDR) were calculated over GO classes, based on P-values from Fisher’s exact tests. Full circles and full lines indicate FDR < 0.05, for predicted and observed PNC respectively.

To analyze the function of genes prioritized by predicted PNC, we estimated their enrichment for GO classes. Significant enrichment was detected for genes involved in catalytic activity and nucleotide binding (e.g., ATP binding for energy transfer). Based on these functional enrichments, predicted PNC prioritized genes involved in primary metabolism (Fig. 5b, Supplementary Fig. S6). In contrast, genes involved in gene regulation and plant development were depleted by these prioritizations. Prioritization by observed PNC also resulted in significant depletion for these GO classes, so PNC across Angiosperms may have de-emphasized developmental genes, possibly because of functional turnover over large evolutionary timescales [21]. Even though we included PNC over moderate evolutionary timescales (tree size between 5 and its maximum, 16.2), clade-specific constraint (e.g., at the genus level) could not be detected in the sample of genomes used in this study [23]. In addition, the depletion by predicted PNC may have been exacerbated by the prediction model itself (Supplementary Fig. S6); the absence of genomic annotations about gene regulation (e.g., RNA-protein binding) may have downplayed the importance of developmental genes for fitness. Finally, these depletions might actually reflect relaxed selection on low gene expression (expression-rate anticorrelation) [23–25]. However, even after accounting for RNA and protein expression, we still observed significant depletions for these GO classes (Supplementary Fig. S6), so we could not rule out genes’ functional importance as a direct determinant of PNC.

### Predicted evolutionary constraint improves genomic prediction in maize

To assess the functional relevance and practical utility of predicted PNC, we used predicted PNC to weight nonsynonymous SNPs in genomic prediction for agronomic traits: days to silking (DTS), plant height (PH) or grain yield (GY). We tested the hypothesis that predicted PNC was larger at causal variants for fitness-related traits in hybrid panels. Under this hypothesis, we expected that (i) weighting SNPs with predicted PNC increased the accuracy of genomic prediction; and (ii) prioritizing SNPs with larger predicted PNC resulted in further gains in accuracy. Expectation (i) was not met for any of the agronomic traits (Supplementary Fig. S7), probably because of the large LD extent in the hybrid panels (average squared correlation above 0.1 within 100-kb distance), such that causal variants were adequately tagged even by randomly weighted SNPs [14].

Expectation (ii) was met for GY, our trait most related to fitness; a gradual increase in prediction accuracy was observed as prioritization of SNPs was more stringent, with a trend similar to that for lower MAF (Supplementary Fig. S4). Moreover, a significant increase was obtained by prioritizing the top 1040 (1%) and 104 (0.1%) SNPs (P < 0.05 based on random permutations of SNP weights). These gains in prediction accuracy were greater than those achieved by observed PNC, despite ∼80 times fewer prioritized SNPs (Fig. 6a). Assuming the minor allele to be deleterious, prioritizations of the top 0.1% SNPs would select 15 mutations per inbred line on average (Table S1). Therefore, stringent prioritization of SNPs by predicted PNC could enable the selection of manageable numbers of candidate variants, for subsequent purging of mutation load by breeding or CRISPR-based editing.

**Figure 6.**
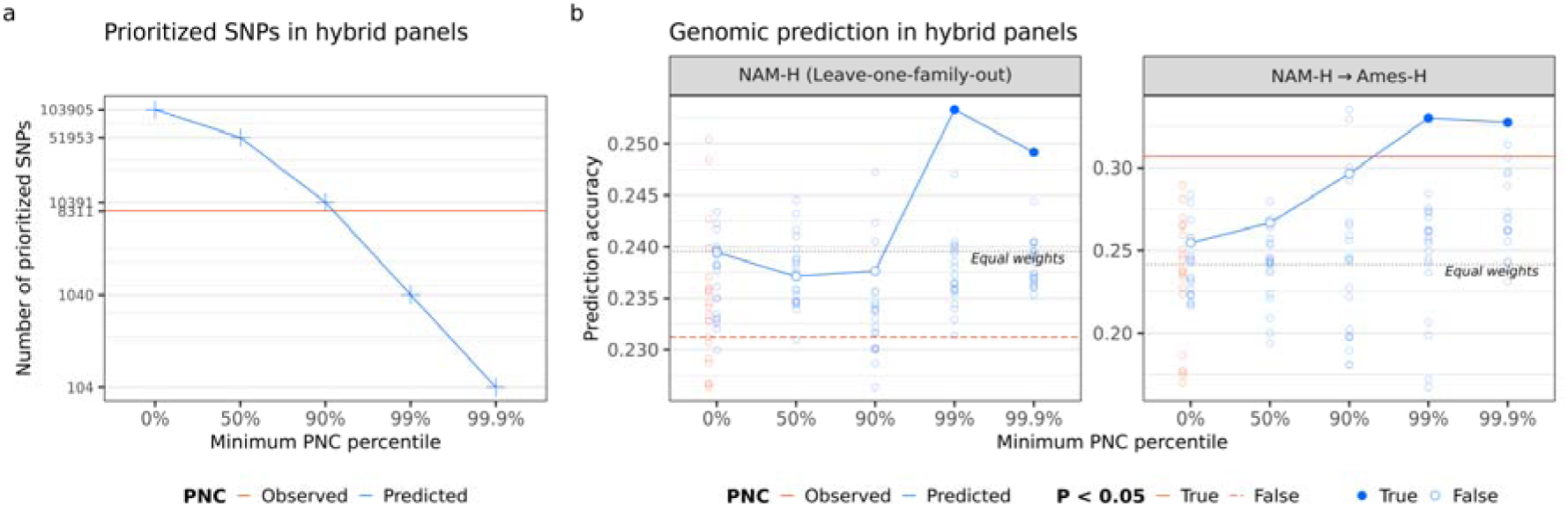
Prioritization of nonsynonymous SNPs in genomic prediction for grain yield, in the Nested Association Mapping hybrid panel (NAM-H). (a) Number of SNPs prioritized by phylogenetic nucleotide conservation (PNC). (b) Genomic prediction accuracy within panel (in leave-one-family-out prediction in NAM-H) or across panels, from NAM-H to a diverse hybrid panel (Ames-H) (14). Black dashed line: nonsynonymous SNPs were weighted equally (“Equal weights”). Red line: nonsynonymous SNPs were weighted by observed PNC. Blue curve: Nonsynonymous SNPs were weighted by predicted PNC, and prioritized by truncating weights to zero if they were under the 0%, 50%, 90%, 99%, or 99.9% quantile. Open circles: nonsynonymous SNPs were weighted and prioritized by random permutations of predicted (blue) or observed (red) PNC. Full circles and full lines indicate P < 0.05 based on random permutations of SNP weights, for predicted and observed PNC respectively. All genomic prediction models accounted for genome-wide effects by principal components and a genomic relationship matrix.

Significant increase in prediction accuracy for GY was observed in a large panel of half-sibs (NAM-H), likely because the effects of deleterious mutations from the recurrent parent were estimated accurately. This gain was significant but modest (0.25 by prioritizing the top 0.1% vs. 0.24 by weighting all nonsynonymous SNPs equally), probably because the donor parents were unrelated and shared few deleterious mutations with one another (Fig. 6b). However, when we used NAM-H to predict GY in a different panel (Ames-H), we achieved a large and significant increase in prediction accuracy (0.33 by prioritizing the top 0.1% vs. 0.24 with equal weights; Fig. 6b). The positive result for GY was not observed when training a genomic prediction model in Ames-H. In this panel representative of maize diversity, variation at SNPs – and the information available to learn their effect – was negatively correlated with species-wide MAF [14]. Therefore, prioritization by PNC of variants with lower MAF (Fig. 4a) resulted in larger estimation errors in Ames-H, and may explain why genomic prediction models trained in this panel benefited less from prioritizations by predicted PNC (Supplementary Fig. S7). Accordingly, enrichment of rare SNPs (MAF < 0.01 in Hapmap 3.2.1) for genomic effects on GY was large and significant in NAM-H (> 16-fold enrichment) but not in Ames-H [14].

Genomic prediction was not improved by PNC for other agronomic traits: PH and DTS. This lack of improvement may be due to a weak relationship between these traits and evolutionary constraint, as proxied by PNC across Angiosperms. Consistently, in maize, hybrid vigor and inbreeding depression are substantially larger for traits related to seed weight and grain yield, compared to traits related to plant morphology and flowering time [28–34]. Interestingly, prioritizations by predicted PNC resulted in a gradual decrease and a significant loss of accuracy for DTS, in a genomic prediction model trained in Ames-H, which suggests that predicted PNC may actually fail to detect variants that are causal for adaptive traits like flowering time (Supplementary Fig. S7). Moreover, enrichment of rare SNPs (MAF < 0.01 in Hapmap 3.2.1) for effects on PH and DTS was not detected in either Ames-H or NAM-H [14], which suggests that the SNPs impacting these traits are under weaker negative selection, compared to those impacting GY. Together, our results on PNC and previous results on MAF indicate that prioritization of SNPs by PNC may improve genomic prediction if causal SNPs are under negative selection, and carry enough statistical information in the training set (e.g., causal SNPs for a yield trait like GY in a collection of biparental populations like NAM-H).

## Discussion

Our results about the characteristics of prioritized SNPs and genes suggest that predicted PNC is more useful than observed PNC to identify causal variants for fitness-related traits, since it can select fewer variants and produce stronger functional enrichments. Our approach (PICNC) addressed two important caveats of observed PNC, which limit its usefulness for quantitative genetics and breeding applications: missing information outside MSAs, and sensitivity of conservation to fitness effects [39, 40]. Our predictions of PNC addressed the first caveat by using, as predictors, genomic annotations that are readily available from genomic and protein sequence data. These genomic annotations were produced by bioinformatic and machine learning procedures which are designed for broad sets of species, with the exception of transposon insertion which was detected by maize-specific transposon motifs [41–43] but was not important in our predictions (Fig. 3b). The second caveat is due to the high sensitivity of PNC, such that PNC tends to its maximum as soon as evolutionary constraint is moderate, especially in MSA across few taxa. Our predictions addressed this caveat by estimating a probability for PNC, which could be used to select arbitrarily small sets of sites in prioritizations, whereas observed PNC might select too many sites for breeding applications like biological design (e.g., 55,789 and 8,311 nonsynonymous SNPs with maximum observed PNC in Hapmap 3.2.1 and hybrid panels, respectively).

Our approach was validated by cross-population genomic prediction (training in one set of populations, validation in a distinct set of populations). Compared to within-population genomic prediction, cross-population genomic prediction accuracies across populations are typically much lower – and sometimes close to zero [36], because of differences in LD patterns and allele frequencies between training set and validation set [34].

Significant improvements in cross-population genomic prediction for GY suggest that prioritization of SNPs by predicted PNC could be useful for breeding applications (e.g., genomic pre-breeding [44], or genomic selection in understudied populations [45]). They also suggest that predicted PNC could point to useful causal variants, because accurate cross-population prediction requires very close tagging of causal variants by genomic markers [46, 47]. Our improvements in prediction accuracy for GY (+5% and +38%) are on par with those achieved from genome-wide prioritization of causal variants, with many experimental annotations in large human samples (trans-ancestry predictions in cohorts of size > 150,000) [48]. However, they suggest that prioritization by PNC is only useful for fitness-related traits, for which causal variants are likely to be under evolutionary constraint. In this study, prioritization by PNC was tested in elite maize populations, in which deleterious mutations have been purged through sustained crop improvement [49]. It could be even more useful in other maize populations [50] or other crop species, in which deleterious mutations are widespread, like sorghum [51] or cassava [11, 51].

Our approach exemplifies important benefits of this coming generation of protein structural machine learning annotations for predicting PNC without resorting to experimental data. It will encourage future studies, which will apply similar approaches to noncoding regulatory variants. Recent studies have introduced promising methods to predict gene regulation and infer high-resolution scores about the effect of mutations, e.g., for TF binding [11, 52], RNA expression [34,53,54] and RNA-protein binding [54]. Our results demonstrate the usefulness of our methodology. They also open possibilities for improved detection of fitness effects, by including broader sets of variants (e.g., noncoding variants), novel genomic annotations (e.g., regulatory effects of genes and mutations) and different evolutionary timescales (e.g., clade-specific fitness effects).

Moreover, further improvements of SNP prioritization could be achieved by combining our approach with complementary techniques. Recent studies in human genetics have inferred relationships between genomic annotations and functional impact of mutations. These include methods based on evolutionary data, like CADD [55] and LINSIGHT [27], as well as methods based on summary statistics from genome-wide association studies (GWAS) [54, 55]. GWAS-based methods are subject to biases from SNP survivorship and LD, but they describe the effect of mutations on explicitly-defined traits. Therefore, these methods could be useful in combination with our proposed method, which does not suffer from the same caveats.

## Conclusions

To detect causal variants at high resolution, we used nucleotide conservation and machine learning to predict the impact of mutations at single DNA sites. Our methodology benefited from genomic annotations which described protein structure by deep neural networks and estimated the structural impact of mutations by *in silico* mutagenesis. In maize, nucleotide conservation predicted by our approach performs better than observed nucleotide conservation. It results in significant functional enrichments, and improves genomic prediction for grain yield across elite populations. Therefore, our approach (PICNC) could enable breeding applications which require the identification of causal variants at high resolution, like cross-population genomic prediction and genome editing.

## Methods

### Training data

#### Genomic data

The B73 maize reference genome and its gene model annotations were downloaded from Ensembl Plants under version 3, release 31 (ftp://ftp.ensemblgenomes.org/pub/plants/release-31/fasta/zea_mays/). Nuclear gene models with 3’UTR and 5’UTR annotations (hereafter, genes) were retained for further analyses (25,824 genes). The representative transcript for each gene model was the transcript with the most matches (bit-score > 50 in global alignment) with any other transcripts in the genomes of B73, Mo17, BTx623 (*Sorghum bicolor*) and Yugu1 (*Setaria italica*), or, by default, the longest transcript. Mutations in the coding region of representative transcripts were characterized at two types of DNA bases: monomorphic sites and SNP sites. Mutations at monomorphic sites were 20,136,310 random nonsynonymous substitutions in the maize genome at the selected genes, while those at SNP sites were the 483,448 observed nonsynonymous substitutions in Hapmap 3.2.1, a representative panel of inbred lines in maize [13].

#### Evolutionary constraint

Publicly available data from a multiple-sequence alignment (MSA) across Angiosperms was previously published in maize [23]: neutral score (depth of MSA at each site) and conservation scores (rejected substitutions) from gerp++ [58]. For each site *j*, phylogenetic nucleotide conservation (PNC) *w_j_* was binary: *w_j_* = 1 f the neutral score (tree size) was > 5 and the ratio of conservation score to neutral score was > 0.95 (i.e., substitution rate < 0.05), *w_j_* = 0 otherwise.

#### Genomic annotations

Each mutation in coding regions was characterized by genomic structure (GC content, *k*-mer frequency and transposon insertion) and protein structure (mutation type, SIFT score, UniRep variables and *in silico* mutagenesis scores).

GC content was the number of G or C bases from -49 to +50 bases from the site of the mutation. *k*-mer frequency was the average frequency of all 13-mers comprising the mutation’s site, calculated by jellyfish [59]. Predictions of transposon insertion at the mutation’s site (helitron, TIR, LINE or LTR) were downloaded from https://github.com/mcstitzer/maize_TEs/blob/master/B73.structuralTEv2.disjoined.2018-09-19.gff3.gz [60].

Mutation type (missense, stop gain or stop loss), SIFT score and SIFT class (“constrained” if SIFT score ≤ 0.05, “tolerated” otherwise) were predicted using SIFT 4G [61]. UniRep variables were the 256 values generated for each protein sequence by the “256-unit UniRep model” available from https://github.com/churchlab/UniRep [61]. *In silico* mutagenesis scores measured the impact of each mutation on proteins, as quantified by the UniRep variables: 256 deviations + 1 Euclidean distance between the reference representation and the mutated representation.

### PICNC: Prediction of evolutionary constraint by genomic annotations

#### Model fitting

The relationship between genomic annotations and observed PNC (*w_j_* = 0 or 1) was estimated by probability random forests [62] implemented in the R package *ranger* [63]. To maximize power to differentiate negative (*w_j_* = 0) and positive examples (*w_j_* = 1) of evolutionary constraint, *w_j_* was set to missing in intermediate cases where substitution rate > 0.005 or tree size < 5 (*w_j_* = 0 only in least conserved regions where the MSA is missing). The probability *P*(*w_j_* = 1) was estimated by 1000 trees per forest, 50,000 sites per tree (sampled with replacement), and at least 100 sites at each terminal node. Mutation effect, SIFT score and SIFT class were always included as baseline predictors, while a third of remaining genomic annotations (GC content, *k*-mer frequency, transposon insertion, UniRep variables and *in silico* mutagenesis scores) were randomly sampled as predictors for each tree. To account for imbalance with respect to PNC and chromosome, each observation (site) was weighted by the inverse of the count of its respective class, as determined by its observed PNC and its chromosome.

#### Leave-one-chromosome-out prediction

For each chromosome *k*: = 1,…,10, PNC at each SNP site in chromosome *k* was predicted by a probability random forest 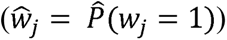, trained on monomorphic sites in all chromosomes except *k* (Fig. 2). Importance of genomic annotations in random forests was estimated by the corrected impurity measure [63]. Classification accuracy was estimated by the percentage of sites for which 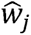 (rounded) equaled *w_j_*, weighted by the sample weights (as described above). When estimating the importance of genomic annotations and assessing the effect of random forest parameters on classification accuracy (number of samples per tree, sets of genomic annotations used in prediction), random forests were validated at monomorphic sites in chromosome 8 and trained (at monomorphic sites) in remaining chromosomes (Fig. 2).

### Validation of predicted evolutionary constraint

#### Experimental SNP annotations

Predicted PNC was validated by measures of functional importance of SNPs: within-species conservation, *cis* eQTL effect, and chromatin accessibility. Within-species conservation was quantified by minor allele frequency (MAF), estimated in a filtered set of SNPs (bi-allelic, minor allele count ≥ 3, missingness ≤ 50%) in the Hapmap 3.2.1 panel [13], imputed by BEAGLE 5.0 [65]. *Cis* eQTL effects were the statistical associations (in absolute value) between SNPs and 3’ RNA-seq expression of genes, in the diverse panel of 299 lines analyzed by [66]. *Cis* eQTL effects in germinating shoot or germinating root were estimated for the SNPs with MAF ≥ 0.05 in this panel, in a linear regression model including the PEER factors from [61] as covariates, using GEMMA 0.98.1 [62]. Chromatin accessibility was characterized by hotspots of MNase hypersensitivity in germinating shoot or germinating root, as defined by [17].

PNC was validated by experimental SNP annotations in a generalized additive model fitted in the R package *mgcv* [63]. PNC was regressed on MAF and *cis* eQTL effects (by cubic regression splines), and chromatin accessibility (as factors), while accounting for chromosome (as factor) and whether the site was included in the MSA (as factor, to control for bias of the MSA towards gene-dense regions).

#### Experimental gene annotations

Predicted PNC was validated by gene properties: gene expression, gene ontology, and ratio of nonsynonymous-to-synonymous SNPs (P_n_/P_s_). Gene expression was quantified by RNA abundance across 23 tissues, and protein abundance across 32 tissues [64]. In all analyses, gene expression was log-transformed: *log*(*x* + 1) where *x* is RNA abundance in Fragments Per Kilobase of transcript per Million mapped reads (FPKM) or protein abundance in distributed normalized spectral abundance factor (dNSAF).

Experimentally-validated gene ontology (GO) annotations [65] were retrieved by mapping protein sequences to the eggNOG database, using DIAMOND [66]. In enrichment analyses, GO annotations were trimmed to the broader (and less redundant) GO slim terms in the “plant GO slim” subset (http://current.geneontology.org/ontology/subsets/goslim_plant.obo), and GO annotations with fewer than 20 positives were discarded (87 selected GO terms). P_n_/P_s_ was the ratio of segregating nonsynonymous SNPs (P_n_) over segregating synonymous SNPs (P_n_) (MAF ≥ 0.01 in Hapmap 3.2.1) within each gene with enough observed segregating synonymous SNPs (Ps ≥ 5).

In validations by experimental gene annotations, genes containing sites with 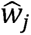 above a threshold value were selected. Threshold values were the 50%, 90%, 99% and 99.9% percentiles of 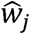’s. Using these successive selections, we assessed the functional enrichment of prioritized genes as fewer sites were included due to more stringent thresholds. The significance of the enrichment for gene expression (difference in mean expression between selected genes and all genes) and GO slim terms (overrepresentation of term among selected genes) were tested by two-sample *t*-test and Fisher’s exact test, respectively.

#### Field traits in hybrid maize

Two panels of hybrid maize lines were analyzed to assess the usefulness of predicted PNC for genomic prediction: a diversity panel (Ames-H; *n*=1106) and a collection of bi-parental crosses having B73 as their common parent (NAM-H; *n*=1640) [14]. These panels were phenotyped for three agronomic traits: days to silking (DTS), plant height (PH) and grain yield adjusted for DTS (GY). They were genotyped for 12,659,487 genome-wide SNPs, including *m*=103,905 nonsynonymous SNPs in the coding regions of the 25,824 genes selected in this study.

Predicted PNC 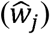 was used to weigh each SNP *j* in genomic prediction models applied to hybrid maize panels:

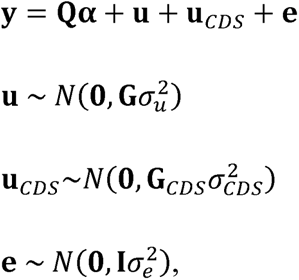

where **y** is the *n*-vector of mean phenotypic values; **Q** is a *n* × 4 matrix depicting population structure by a column of ones (for the intercept) and the three principal components from the Hapmap 3.2.1 panel, with respective effects ; **e** is the vector of **α**; **e** is the vector of errors; **G** is the *n* × *n* genome-wide relationship matrix such that the *n*-vector **u** consists of genome-wide breeding values:

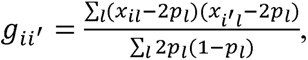

where *x_il_* is the genotype of hybrid *i* at SNP *l*, *p_l_* is the estimated frequency of SNP *l* in hybrid panels.

**G***_CDS_* is the *n* × *n* relationship matrix from nonsynonymous SNPs weighted by predicted PNC, such that the *n*-vector **u***_CDS_* consists of breeding values due to weighted nonsynonymous SNPs:

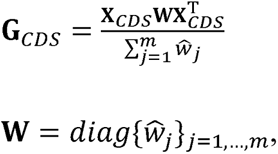

where **X***_CDS_* is the *n* × *m* matrix of genotypes at nonsynonymous SNPs.

Genomic prediction models were fitted by REML, using the R package *regress* [67]. Genomic prediction accuracy was estimated by the Pearson correlation between predicted and observed phenotypic values:

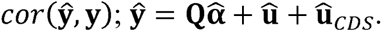

In validations of predicted PNC by genomic prediction, 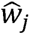’s below a threshold value were set to zero. Threshold values were the 0%, 50%, 90%, 99% and 99.9% percentiles of 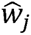’s, among the *m* SNPs observed in hybrid panels. Using these successive truncations, we assessed the enrichment of prioritized SNPs for genomic prediction accuracy, as fewer of them were included due to more stringent thresholding on their weights. The significance of 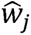’s as useful weights in genomic prediction was tested by comparing genomic prediction accuracy with the accuracies achieved by 20 random permutations of 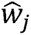’s, hence testing the null hypothesis that 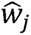’s are as useful as expected by chance.

---

**Figure S1.**
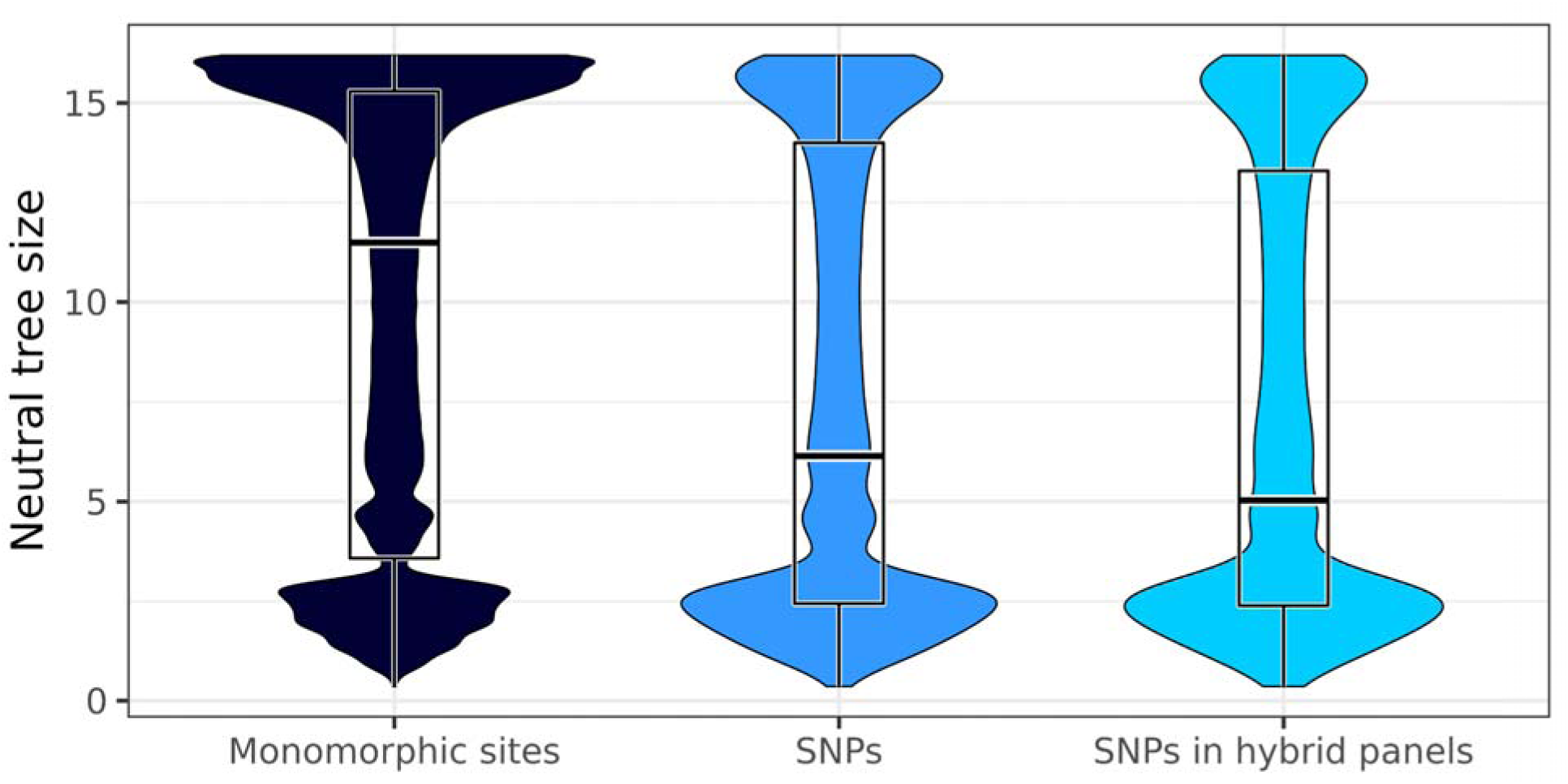
Distribution of neutral tree size (neutral scores) by category of DNA bases. Neutral tree size was calculated as previously described [23]. Monomorphic sites: sites with no observed polymorphism within maize. SNPs: observed polymorphisms in Hapmap 3.2.1, a representative panel of inbred lines in maize [13]. SNPs in hybrid panels: subset of SNPs observed in Hapmap 3.2.1, which are also observed in two panels of hybrid crosses between inbred lines and testers [14].

**Figure S2.**
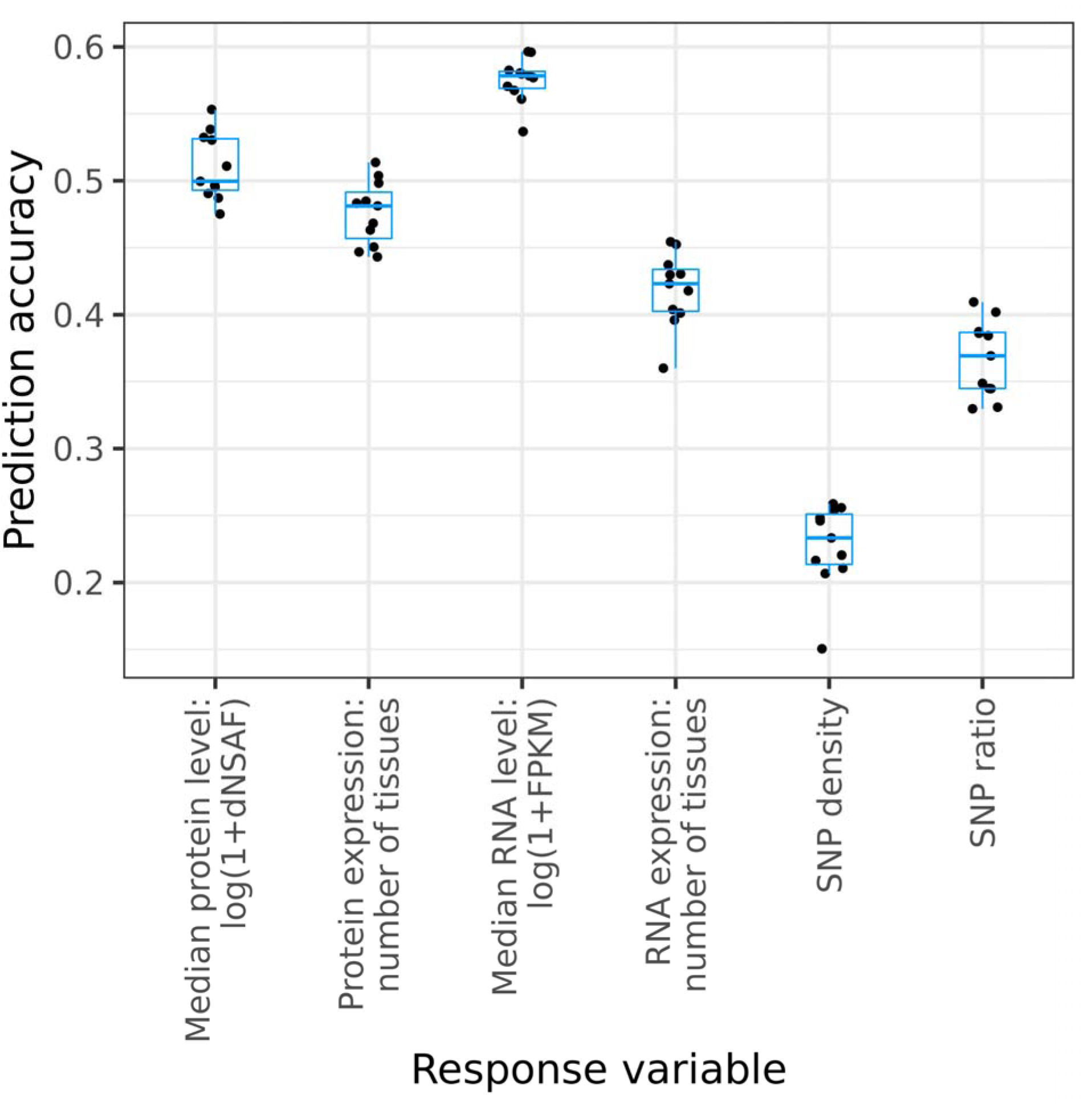
Prediction accuracy of probability random forests for experimental gene annotations, by UniRep variables. UniRep variables are generated by the 256-unit UniRep model. Prediction accuracy is the Pearson correlation coefficient between predicted values and observed values. Expression is quantified by RNA abundance (over 23 tissues) and protein abundance (over 32 tissues) based on the gene expression atlas of [64]: median expression, and number of tissues with non-zero expression level. SNP density: percentage of segregating SNP sites in genes (MAF ≥ 0.01 in Hapmap 3.2.1). SNP ratio: ratio of nonsynonymous-to-synonymous SNPs (P_n_/P_s_) within each gene.

**Figure S3.**
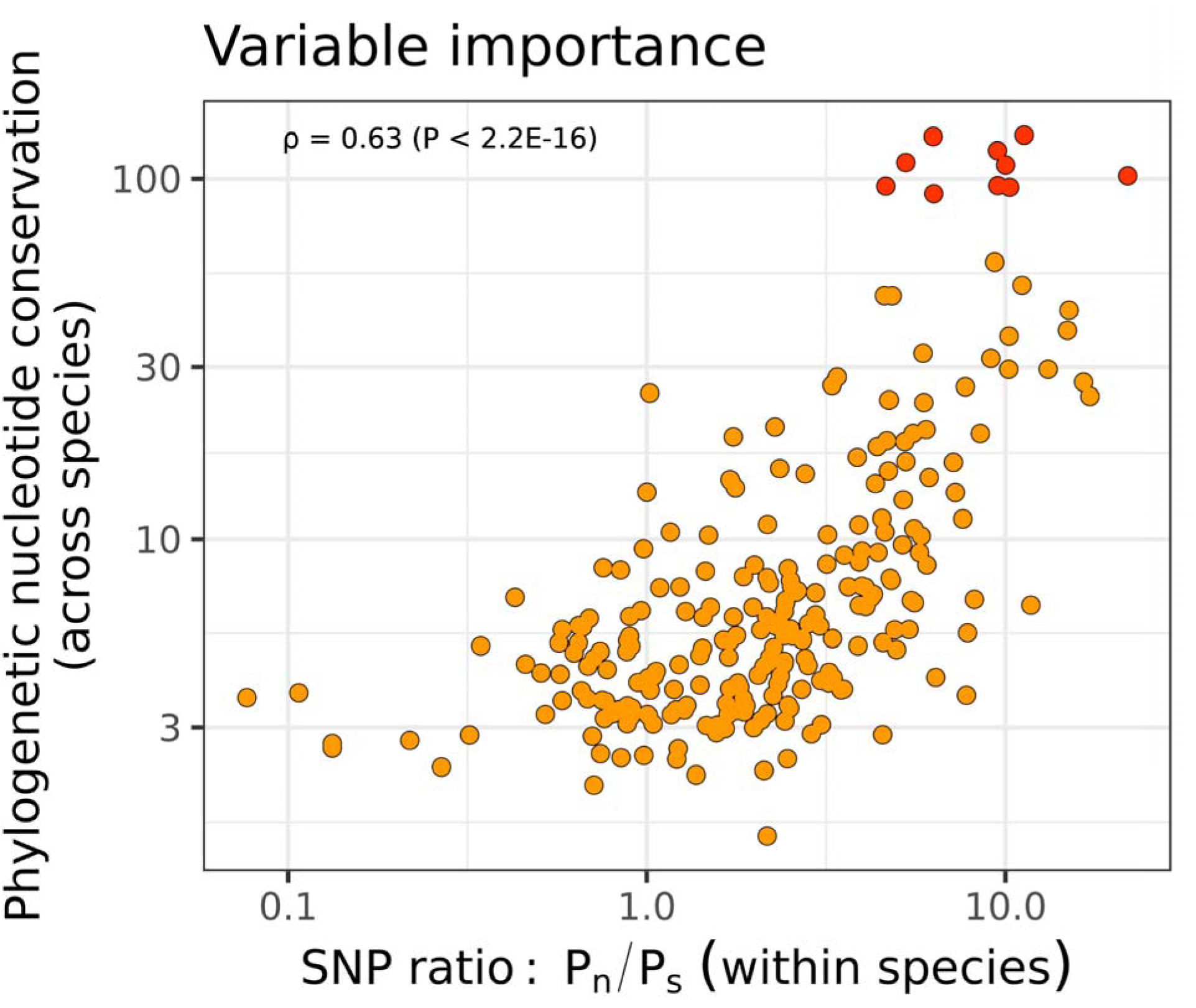
Concordance between importance of UniRep variables for phylogenetic nucleotide conservation (PNC) across species and the ratio of nonsynonymous-to-synonymous SNPs (P_n_/P_s_) within species. The importance of UniRep variables for PNC is correlated with their importance for P_n_/P_s_ at maize genes; ρ: Spearman correlation coefficient.

**Figure S4.**
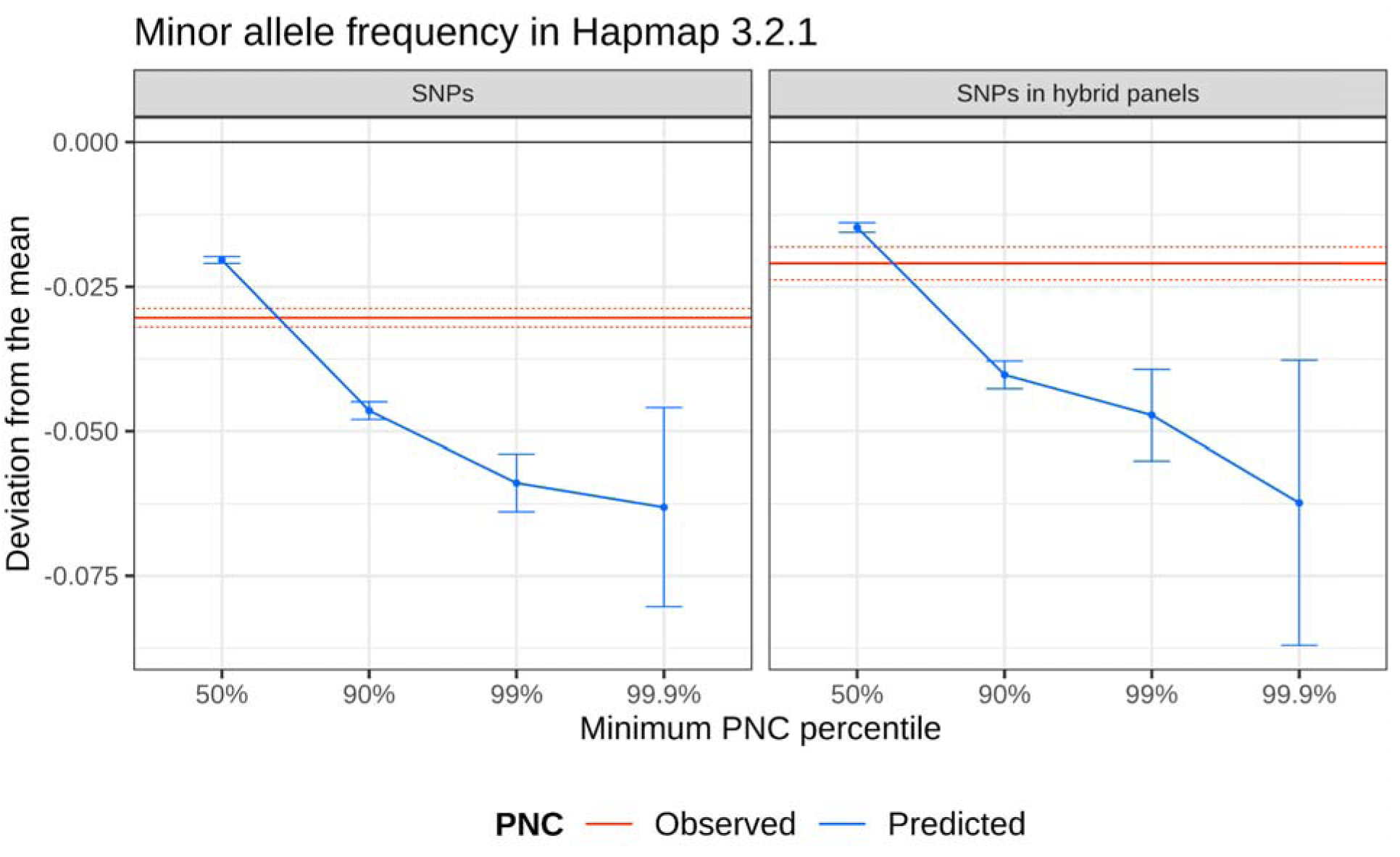
Decrease in minor allele frequency of SNPs prioritized by phylogenetic nucleotide conservation (PNC). Difference in minor allele frequency between prioritized SNPs and all SNPs. SNPs: observed polymorphisms in Hapmap 3.2.1, a representative panel of inbred lines in maize [13]. SNPs in hybrid panels: subset of SNPs observed in Hapmap 3.2.1, which are also observed in two panels of hybrid crosses between inbred lines and testers [14]. SNPs were prioritized if their predicted PNC was above the 50%, 90%, 99%, or 99.9% quantile, or their observed PNC was equal to 1 (tree size > 5, substitution rate < 0.05). Error bars and dotted lines represent 95% confidence intervals in two-sample t-tests, for predicted and observed PNC respectively.

**Figure S5.**
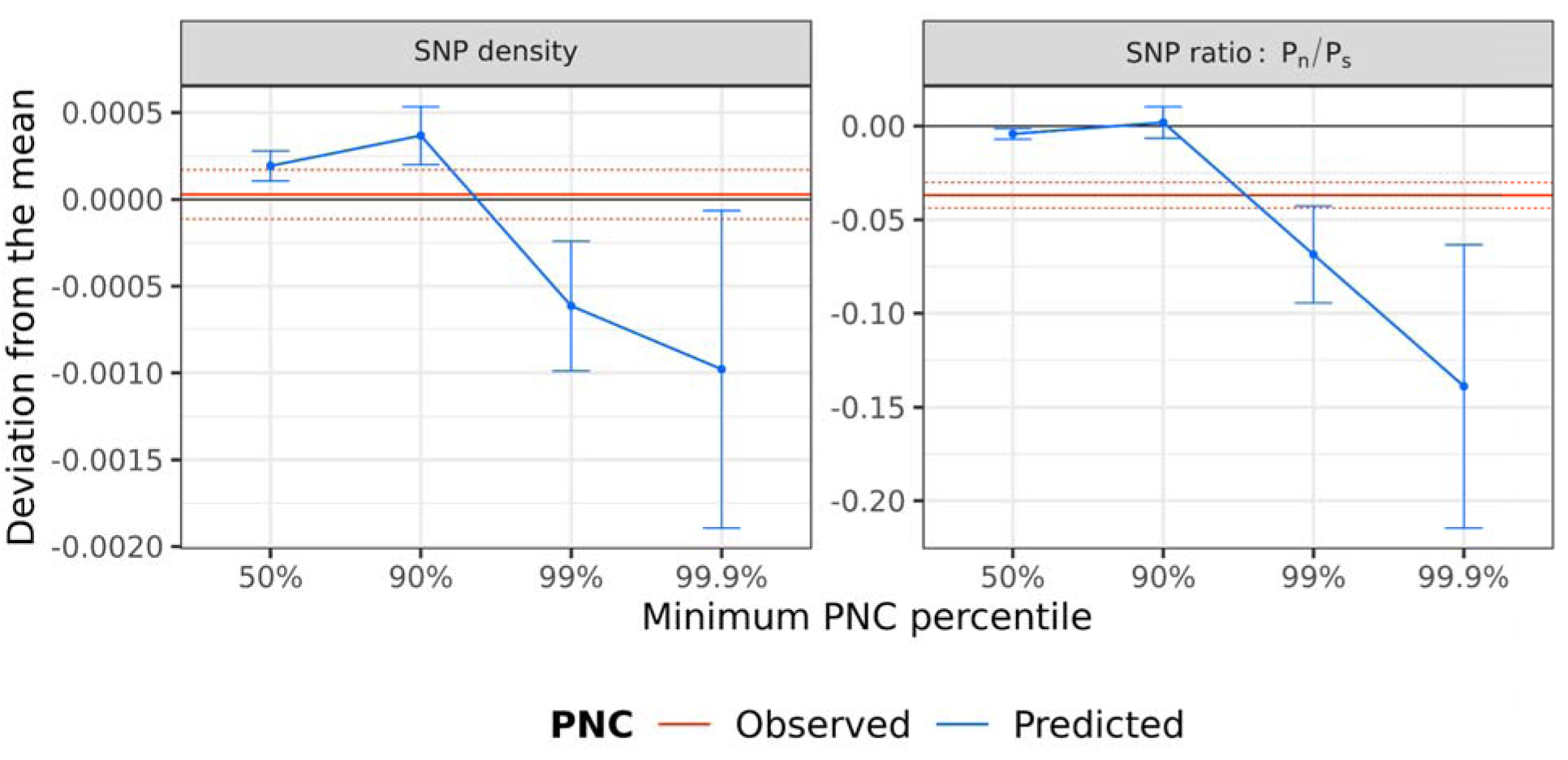
Difference in experimental annotations of genes prioritized by phylogenetic nucleotide conservation (PNC). Difference in experimental annotations between prioritized genes and all genes. SNP density: percentage of sites for which a SNP is observed (MAF ≥ 0.01 in Hapmap 3.2.1) within each gene. SNP ratio: ratio of nonsynonymous-to-synonymous SNPs (P_n_/P_s_) within each gene. Genes were prioritized by selecting SNPs with a predicted PNC above the 50%, 90%, 99%, or 99.9% quantile, or observed PNC equal to 1 (tree size > 5, substitution rate < 0.05). Error bars and dotted lines represent 95% confidence intervals in two-sample t-tests, for predicted and observed PNC respectively.

**Figure S6.**
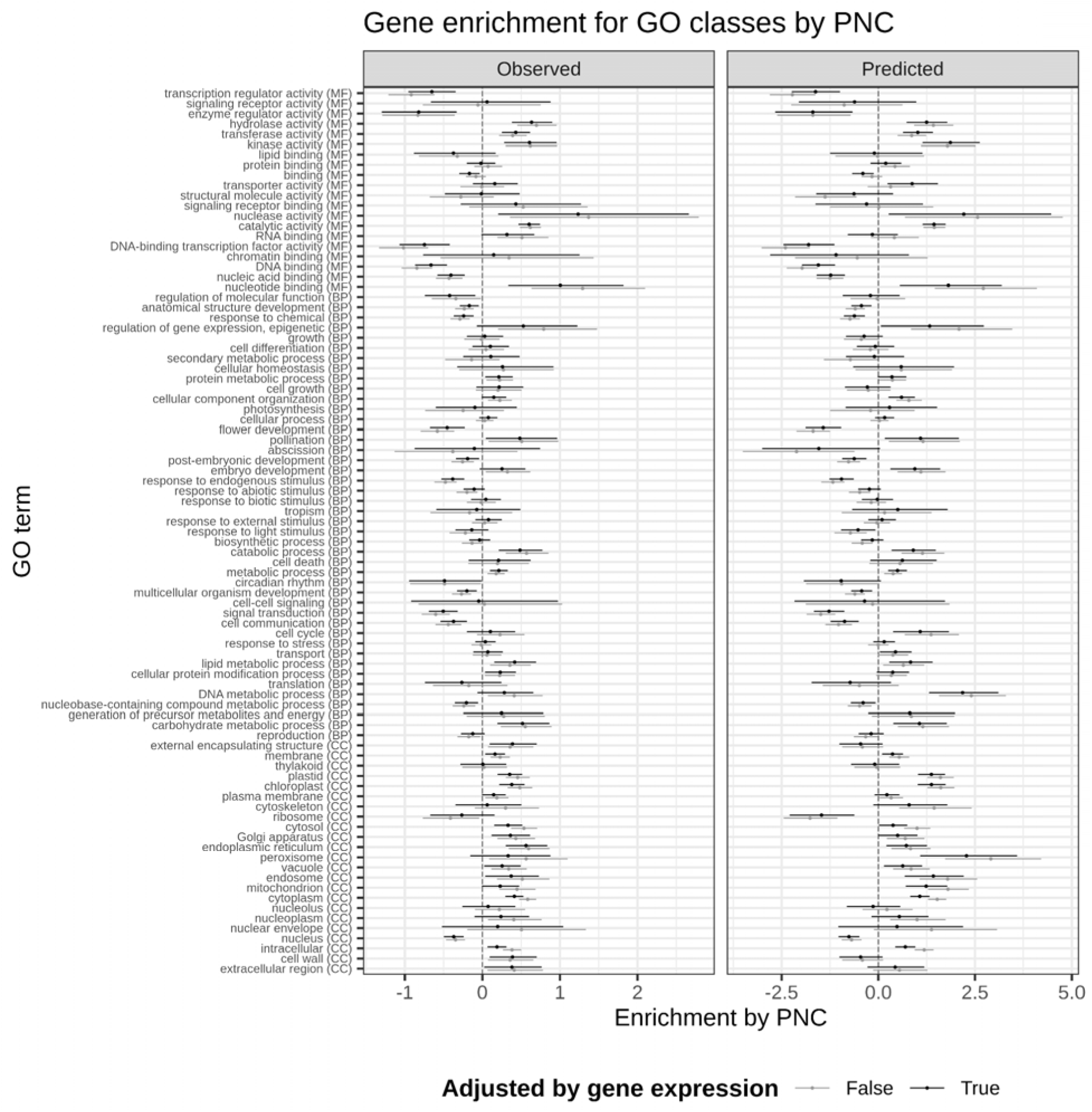
Enrichment of genes prioritized by phylogenetic nucleotide conservation (PNC), for gene ontology (GO) classes. Effect of maximum PNC (in each gene coding sequence) on the odds ratio for GO annotations [Pr(GO annotation)/(1 - Pr(GO annotation)], based on logistic regression. Estimated effects of gene prioritizations are shown on a log scale: point estimates (dot) and 95%-confidence intervals (segment). Gray symbols: effects of gene prioritizations are not adjusted (simple logistic regression of GO annotation on maximum PNC). Black symbols: effects of gene prioritizations are adjusted by gene expression (logistic regression including gene expression variables as covariates). Gene expression variables are RNA abundance (FPKM over 23 tissues) and protein abundance (dNSAF over 32 tissues) based on the gene expression atlas of [64]: median expression [median of log(1+FPKM) or log(1+dNSAF)], and number of tissues with non-zero expression level. GO classes belong to the plant GO slim subset. Ontology: CC, cellular component; BP, biological process; MF: molecular function.

**Figure S7.**
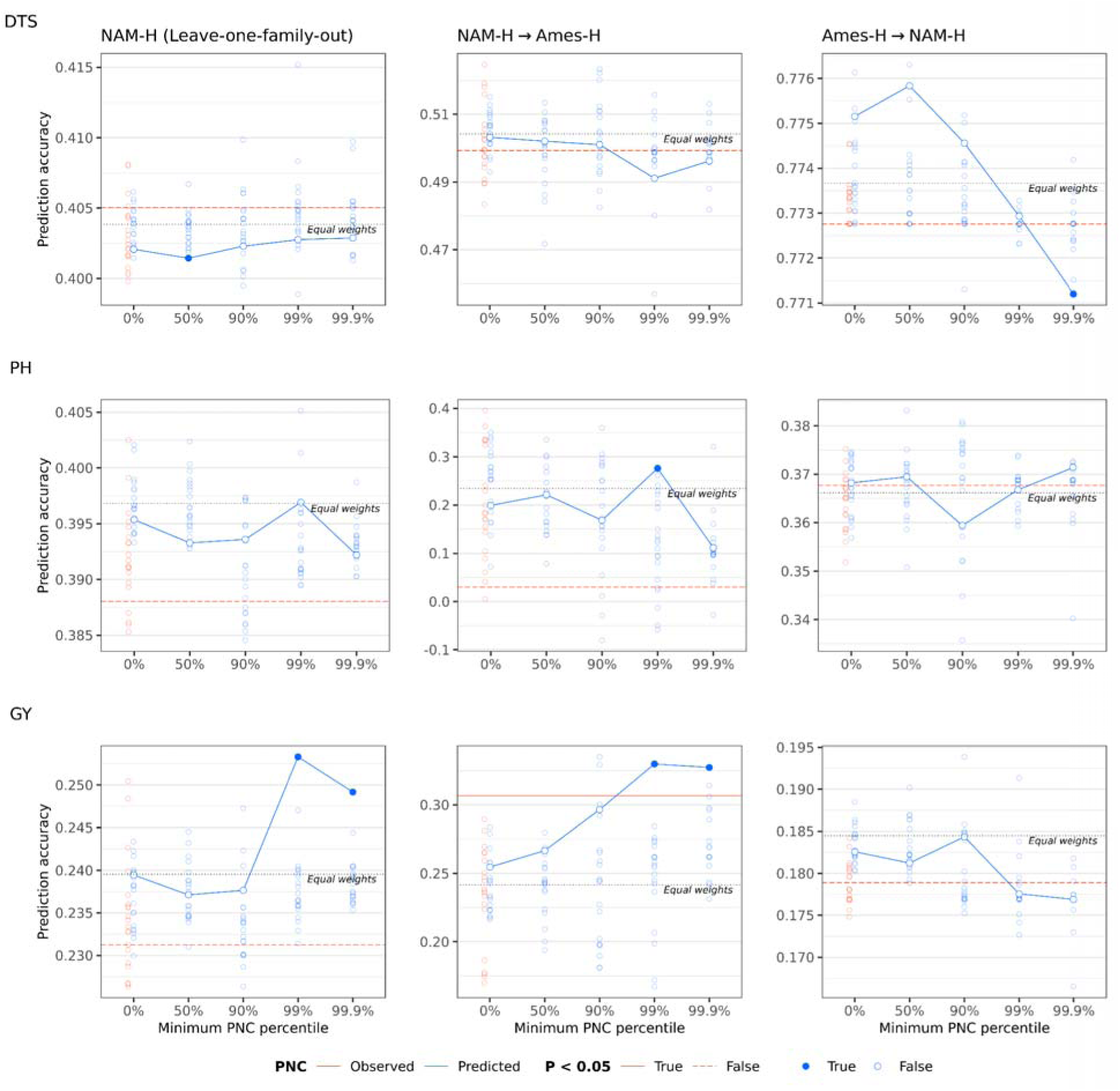
Prioritization of nonsynonymous SNPs in genomic prediction for agronomic traits in hybrid panels. Agronomic traits: days to silking (DTS), plant height (PH) and grain yield (GY); hybrid panels: Nested Association Mapping hybrid panel (NAM-H), diverse hybrid panel (Ames-H) [14]. Genomic prediction accuracy was estimated within NAM-H (in leave-one-family-out prediction), from NAM-H to Ames-H, and from Ames-H to NAM-H. Black dashed line: nonsynonymous SNPs were weighted equally (“Equal weights”). Red line: nonsynonymous SNPs were weighted by observed phylogenetic nucleotide conservation (PNC). Blue curve: nonsynonymous SNPs were weighted by predicted PNC, and prioritized by truncating weights to zero if they were under the 0%, 50%, 90%, 99%, or 99.9% quantile. Open circles: nonsynonymous SNPs were weighted and prioritized by random permutations of predicted (blue) or observed (red) PNC. Full circles and full lines indicate P < 0.05 based on random permutations of SNP weights, for predicted and observed PNC respectively. All genomic prediction models accounted for genome-wide effects by principal components and a genomic relationship matrix.

**Table S1.** Prioritization of nonsynonymous SNPs by phylogenetic nucleotide conservation (PNC): number of selected SNPs in hybrid panels and expected number of deleterious mutations per inbred lines based on minor allele frequency (MAF) in Hapmap 3.2.1

| **PNC** | **Minimum PNC percentile (%)** | **Number of prioritized SNPs** | **Average MAF in Hapmap 3.2.1** | **Number of minor alleles per haploid genome** |
| --- | --- | --- | --- | --- |
| **Observed** | NA | 8311 | 0.185 | 1537.2 |
| **Predicted** | 0 | 103905 | 0.206 | 21394.3 |
|  | 50 | 51953 | 0.191 | 9933.5 |
|  | 90 | 10391 | 0.166 | 1721.4 |
|  | 99 | 1040 | 0.159 | 165.1 |
|  | 99.9 | 104 | 0.144 | 14.9 |

## References

1. Tam V, Patel N, Turcotte M, Bossé Y, Paré G, Meyre D. Benefits and limitations of genome-wide association studies. Nat Rev Genet. nature.com; 2019;20:467–84.

2. Lanfear R, Kokko H, Eyre-Walker A. Population size and the rate of evolution. Trends Ecol Evol. 2014;29:33–41.

3. Vaser R, Adusumalli S, Leng SN, Sikic M, Ng PC. SIFT missense predictions for genomes. Nat Protoc. nature.com; 2016;11:1–9.

4. Davydov EV, Goode DL, Sirota M, Cooper GM, Sidow A, Batzoglou S. Identifying a high fraction of the human genome to be under selective constraint using GERP++. PLoS Comput Biol. 2010;6:e1001025.

5. Rands CM, Meader S, Ponting CP, Lunter G. 8.2% of the Human genome is constrained: variation in rates of turnover across functional element classes in the human lineage. PLoS Genet. 2014;10:e1004525.

6. Huber CD, Kim BY, Lohmueller KE. Population genetic models of GERP scores suggest pervasive turnover of constrained sites across mammalian evolution. PLoS Genet. journals.plos.org; 2020;16:e1008827.

7. Kimura M. On the probability of fixation of mutant genes in a population. Genetics. 1962;47:713–9.

8. Kircher M, Witten DM, Jain P, O’Roak BJ, Cooper GM, Shendure J. A general framework for estimating the relative pathogenicity of human genetic variants. Nat Genet. 2014;46:310–5.

9. Rentzsch P, Witten D, Cooper GM, Shendure J, Kircher M. CADD: predicting the deleteriousness of variants throughout the human genome. Nucleic Acids Res. academic.oup.com; 2019;47:D886–94.

10. Arbiza L, Gronau I, Aksoy BA, Hubisz MJ, Gulko B, Keinan A, et al. Genome-wide inference of natural selection on human transcription factor binding sites. Nat Genet. nature.com; 2013;45:723–9.

11. Huang Y-F, Gulko B, Siepel A. Fast, scalable prediction of deleterious noncoding variants from functional and population genomic data. Nat Genet. 2017;49:618–24.

12. Alley EC, Khimulya G, Biswas S, AlQuraishi M, Church GM. Unified rational protein engineering with sequence-based deep representation learning. Nat Methods. 2019;16:1315–22.

13. Bukowski R, Guo X, Lu Y, Zou C, He B, Rong Z, et al. Construction of the third-generation Zea mays haplotype map. Gigascience. academic.oup.com; 2018;7:1–12.

14. Ramstein GP, Larsson SJ, Cook JP, Edwards JW, Ersoz ES, Flint-Garcia S, et al. Dominance Effects and Functional Enrichments Improve Prediction of Agronomic Traits in Hybrid Maize. Genetics. Genetics Soc America; 2020;215:215–30.

15. Bierne N, Eyre-Walker A. The genomic rate of adaptive amino acid substitution in Drosophila. Mol Biol Evol. 2004;21:1350–60.

16. Mezmouk S, Ross-Ibarra J. The pattern and distribution of deleterious mutations in maize. G3 . g3journal.org; 2014;4:163–71.

17. Rodgers-Melnick E, Vera DL, Bass HW, Buckler ES. Open chromatin reveals the functional maize genome. Proc Natl Acad Sci U S A. 2016;113:E3177–84.

18. Pál C, Papp B, Hurst LD. Highly expressed genes in yeast evolve slowly. Genetics. academic.oup.com; 2001;158:927–31.

19. Drummond DA, Bloom JD, Adami C, Wilke CO, Arnold FH. Why highly expressed proteins evolve slowly. Proc Natl Acad Sci U S A. National Acad Sciences; 2005;102:14338–43.

20. Yang J-R, Liao B-Y, Zhuang S-M, Zhang J. Protein misinteraction avoidance causes highly expressed proteins to evolve slowly. Proc Natl Acad Sci U S A. National Acad Sciences; 2012;109:E831–40.

21. Park C, Chen X, Yang J-R, Zhang J. Differential requirements for mRNA folding partially explain why highly expressed proteins evolve slowly. Proc Natl Acad Sci U S A. National Acad Sciences; 2013;110:E678–86.

22. Zhang J, Yang J-R. Determinants of the rate of protein sequence evolution. Nat Rev Genet. 2015;16:409–20.

23. Kistler L, Maezumi SY, Gregorio de Souza J, Przelomska NAS, Malaquias Costa F, Smith O, et al. Multiproxy evidence highlights a complex evolutionary legacy of maize in South America. Science. science.sciencemag.org; 2018;362:1309–13.

24. Springer NM, Stupar RM. Allelic variation and heterosis in maize: how do two halves make more than a whole? Genome Res. genome.cshlp.org; 2007;17:264–75.

25. Flint-Garcia SA, Buckler ES, Tiffin P, Ersoz E, Springer NM. Heterosis is prevalent for multiple traits in diverse maize germplasm. PLoS One. 2009;4:e7433.

26. Larièpe A, Mangin B, Jasson S, Combes V, Dumas F, Jamin P, et al. The genetic basis of heterosis: multiparental quantitative trait loci mapping reveals contrasted levels of apparent overdominance among traits of agronomical interest in maize (Zea mays L.). Genetics. Genetics Soc America; 2012;190:795–811.

27. Stitzer MC, Anderson SN, Springer NM, Ross-Ibarra J. The genomic ecosystem of transposable elements in maize. PLoS Genet. 2021;17:e1009768.

28. Lehermeier C, Krämer N, Bauer E, Bauland C, Camisan C, Campo L, et al. Usefulness of multiparental populations of maize (Zea mays L.) for genome-based prediction. Genetics. 2014;198:3–16.

29. Ramstein GP, Casler MD. Extensions of BLUP Models for Genomic Prediction in Heterogeneous Populations: Application in a Diverse Switchgrass Sample. G3 . 2019;9:789–805.

30. Juliana P, Singh RP, Poland J, Mondal S, Crossa J, Montesinos-López OA, et al. Prospects and Challenges of Applied Genomic Selection-A New Paradigm in Breeding for Grain Yield in Bread Wheat. Plant Genome [Internet]. 2018;11. Available from: http://dx.doi.org/10.3835/plantgenome2018.03.0017

31. Kachman SD, Spangler ML, Bennett GL, Hanford KJ, Kuehn LA, Snelling WM, et al. Comparison of molecular breeding values based on within- and across-breed training in beef cattle. Genet Sel Evol. Springer; 2013;45:30.

32. Raymond B, Bouwman AC, Schrooten C, Houwing-Duistermaat J, Veerkamp RF. Utility of whole-genome sequence data for across-breed genomic prediction. Genet Sel Evol. 2018;50:27.

33. Martin AR, Gignoux CR, Walters RK, Wojcik GL, Neale BM, Gravel S, et al. Human Demographic History Impacts Genetic Risk Prediction across Diverse Populations. Am J Hum Genet. 2017;100:635–49.

34. Amariuta T, Ishigaki K, Sugishita H, Ohta T, Koido M, Dey KK, et al. Improving the trans-ancestry portability of polygenic risk scores by prioritizing variants in predicted cell-type-specific regulatory elements. Nat Genet. 2020;52:1346–54.

35. Wientjes YCJ, Veerkamp RF, Calus MPL. Using selection index theory to estimate consistency of multi-locus linkage disequilibrium across populations. BMC Genet. 2015;16:87.

36. van den Berg I, Boichard D, Guldbrandtsen B, Lund MS. Using Sequence Variants in Linkage Disequilibrium with Causative Mutations to Improve Across-Breed Prediction in Dairy Cattle: A Simulation Study. G3 . academic.oup.com; 2016;6:2553–61.

37. Scutari M, Mackay I, Balding D. Using Genetic Distance to Infer the Accuracy of Genomic Prediction. PLoS Genet. 2016;12:e1006288.

38. Cavazos TB, Witte JS. Inclusion of variants discovered from diverse populations improves polygenic risk score transferability. Human Genetics and Genomics Advances. 2021;2:100017.

39. Milner SG, Jost M, Taketa S, Mazón ER, Himmelbach A, Oppermann M, et al. Genebank genomics highlights the diversity of a global barley collection. Nat Genet. 2019;51:319–26.

40. Mascher M, Schreiber M, Scholz U, Graner A, Reif JC, Stein N. Genebank genomics bridges the gap between the conservation of crop diversity and plant breeding. Nat Genet. 2019;51:1076–81.

41. Crossa J, Jarquín D, Franco J, Pérez-Rodríguez P, Burgueño J, Saint-Pierre C, et al. Genomic Prediction of Gene Bank Wheat Landraces. G3 . 2016;6:1819–34.

42. Yu X, Li X, Guo T, Zhu C, Wu Y, Mitchell SE, et al. Genomic prediction contributing to a promising global strategy to turbocharge gene banks. Nat Plants. 2016;2:16150.

43. Dzievit MJ, Guo T, Li X, Yu J. Comprehensive analytical and empirical evaluation of genomic prediction across diverse accessions in maize. Plant Genome. 2021;14:e20160.

44. Chia J-M, Song C, Bradbury PJ, Costich D, de Leon N, Doebley J, et al. Maize HapMap2 identifies extant variation from a genome in flux. Nat Genet. nature.com; 2012;44:803–7.

45. Wang L, Beissinger TM, Lorant A, Ross-Ibarra C, Ross-Ibarra J, Hufford MB. The interplay of demography and selection during maize domestication and expansion. Genome Biol. genomebiology.biomedcentral.com; 2017;18:215.

46. Valluru R, Gazave EE, Fernandes SB, Ferguson JN, Lozano R, Hirannaiah P, et al. Deleterious Mutation Burden and Its Association with Complex Traits in Sorghum (Sorghum bicolor). Genetics. 2019;211:1075–87.

47. Lozano R, Gazave E, Dos Santos JPR, Stetter MG, Valluru R, Bandillo N, et al. Comparative evolutionary genetics of deleterious load in sorghum and maize. Nat Plants. nature.com; 2021;7:17–24.

48. Ramu P, Esuma W, Kawuki R, Rabbi IY, Egesi C, Bredeson JV, et al. Cassava haplotype map highlights fixation of deleterious mutations during clonal propagation. Nat Genet. 2017;49:959–63.

49. Avsec Ž, Weilert M, Shrikumar A, Krueger S, Alexandari A, Dalal K, et al. Base-resolution models of transcription-factor binding reveal soft motif syntax. Nat Genet. 2021;53:354–66.

50. Zhou J, Theesfeld CL, Yao K, Chen KM, Wong AK, Troyanskaya OG. Deep learning sequence-based ab initio prediction of variant effects on expression and disease risk. Nat Genet. nature.com; 2018;50:1171–9.

51. Su Y, Luo Y, Zhao X, Liu Y, Peng J. Integrating thermodynamic and sequence contexts improves protein-RNA binding prediction. PLoS Comput Biol. 2019;15:e1007283.

52. Gronau I, Arbiza L, Mohammed J, Siepel A. Inference of natural selection from interspersed genomic elements based on polymorphism and divergence. Mol Biol Evol. academic.oup.com; 2013;30:1159–71.

53. Gazal S, Loh P-R, Finucane HK, Ganna A, Schoech A, Sunyaev S, et al. Functional architecture of low-frequency variants highlights strength of negative selection across coding and non-coding annotations. Nat Genet. 2018;50:1600–7.

54. Speed D, Holmes J, Balding DJ. Evaluating and improving heritability models using summary statistics. Nat Genet. nature.com; 2020;52:458–62.

55. Marçais G, Kingsford C. A fast, lock-free approach for efficient parallel counting of occurrences of k-mers. Bioinformatics. 2011;27:764–70.

56. Breiman L. Random Forests. Mach Learn. Springer; 2001;45:5–32.

57. Malley JD, Kruppa J, Dasgupta A, Malley KG, Ziegler A. Probability machines: consistent probability estimation using nonparametric learning machines. Methods Inf Med. ncbi.nlm.nih.gov; 2012;51:74–81.

58. Wright MN, Ziegler A. ranger: A Fast Implementation of Random Forests for High Dimensional Data in C++ and R [Internet]. arXiv [stat.ML]. 2015. Available from: http://arxiv.org/abs/1508.04409

59. Nembrini S, König IR, Wright MN. The revival of the Gini importance? Bioinformatics. Oxford University Press (OUP); 2018;34:3711–8.

60. Browning BL, Zhou Y, Browning SR. A One-Penny Imputed Genome from Next-Generation Reference Panels. Am J Hum Genet. Elsevier; 2018;103:338–48.

61. Kremling KAG, Chen S-Y, Su M-H, Lepak NK, Romay MC, Swarts KL, et al. Dysregulation of expression correlates with rare-allele burden and fitness loss in maize. Nature. 2018;555:520–3.

62. Zhou X, Stephens M. Genome-wide efficient mixed-model analysis for association studies. Nat Genet. 2012;44:821–4.

63. Wood SN. Fast stable restricted maximum likelihood and marginal likelihood estimation of semiparametric generalized linear models. J R Stat Soc Series B Stat Methodol. Wiley Online Library; 2011;73:3–36.

64. Walley JW, Sartor RC, Shen Z, Schmitz RJ, Wu KJ, Urich MA, et al. Integration of omic networks in a developmental atlas of maize. Science. 2016;353:814–8.

65. Ashburner M, Ball CA, Blake JA, Botstein D, Butler H, Michael Cherry J, et al. Gene Ontology: tool for the unification of biology. Nat Genet. Nature Publishing Group; 2000;25:25–9.

66. Huerta-Cepas J, Forslund K, Coelho LP, Szklarczyk D, Jensen LJ, von Mering C, et al. Fast Genome-Wide Functional Annotation through Orthology Assignment by eggNOG-Mapper. Mol Biol Evol. 2017;34:2115–22.

67. Clifford D, McCullagh P. The regress function. The Newsletter of the R Project Volume 6/2, May 2006. stat.uchicago.edu; 2005;39243:6.

